# The high-throughput gene prediction of more than 1,700 eukaryote genomes using the software package EukMetaSanity

**DOI:** 10.1101/2021.07.25.453296

**Authors:** Christopher J. Neely, Sarah K. Hu, Harriet Alexander, Benjamin J. Tully

## Abstract

Gene prediction and annotation for eukaryotic genomes is challenging with large data demands and complex computational requirements. For most eukaryotes, genomes are recovered from specific target taxa. However, it is now feasible to reconstruct or sequence hundreds of metagenome-assembled genomes (MAGs) or single-amplified genomes directly from the environment. To meet this forth-coming wave of eukaryotic genome generation, we introduce EukMetaSanity, which combines state-of-the-art tools into three pipelines that have been specifically designed for extensive parallelization on high-performance computing infrastructure. EukMetaSanity performs an automated taxonomy search against a protein database of 1,482 species to identify phylogenetically compatible proteins to be used in downstream gene prediction. We present the results for intron, exon, and gene locus prediction for 112 genomes collected from NCBI, including fungi, plants, and animals, along with 1,669 MAGs and demonstrate that EukMetaSanity can provide reliable preliminary gene predictions for a single target taxon or at scale for hundreds of MAGs. EukMetaSanity is freely available at https://github.com/cjneely10/EukMetaSanity.

## Main

Until recently, the large-scale annotation of eukaryotic genomes has not been a major requirement or consideration for the tools and pipelines built to perform aspects of gene prediction. This is a logical status of the state of the field as standard eukaryotic genomics requires a target organism of interest and extensive financial and data investments for sequencing, both for chromosome construction *i.e*., DNA-centric) and gene locus prediction (*i.e*., RNA-centric) (Xu et al., 2020; Mock et al.,2017; Shoguchi et al., 2018; Li et al., 2018; Leclère et al., 2019). However, with the maturation of the large-scale recovery of eukaryotic metagenomic-assembled genomes (MAGs) (Alexander et al.,2021; Delmont et al., 2021; Duncan et al., 2020; West et al., 2018), the steps for accurately predicting gene loci needs to shift from current methods that typically focus on a single genome to an approach that can be readily parallelized to annotate hundreds or thousands of genomes.

As techniques such as metagenomics and single-cell amplified genomics more frequently provide environmental eukaryotic genomes without the presence of accompanying expression data, accurate gene identification using *ab initio* predictions and protein evidence will be required to leverage the information stored therein. Without the aid of expression data, gene locus prediction is computationally complex and requires two major steps: (1) repeat identification/masking and (2) exon-intron boundary identification (Faure et al., 2021; Salzberg, 2019; Danchin et al., 2018; Yandell and Ence,2012). Both of these steps can be performed by a number of tools/pipelines built to support specific tasks, each with nuances in runtime, input requirements, and user supervision (Bruna et al., 2020; Lomsadze et al., 2005, 2014; Stanke et al., 2006; Hoff and Stanke, 2013; Hoff et al., 2015; Bruna et al., 2021; Levy Karin et al., 2020; Holt and Yandell, 2011; Cantarel et al., 2008). These tasks can also be expedited if suitably close phylogenetic neighbors can be identified to assist in evidence supported prediction(s) (West et al., 2018). Execution times for these steps in a typical eukaryotic annotation pipeline can add hours or days to the total runtime needed to properly identify genes, making large-scale annotation projects difficult to plan and manage.

Here, we present EukMetaSanity, a workflow package that combines crucial steps for accurate gene loci prediction without gene expression data, while also providing downstream avenues for protein annotation and gene refinement through the use of expression data when available (RNA-seq or transcriptomes). EukMetaSanity combines a number of tools and tasks into a single unified package that is easily deployable in different compute environments. The flexibility of EukMetaSanity is built into the application programming interface (API) that allows end-users to select the tools and databases relevant to their questions, but also for rapid incorporation of new tools and high-level parallelization on high performance computing (HPC) systems that support Simple Linux Utility for Resource Management (SLURM) (Yoo et al., 2003).

The overarching workflow for genome annotation is split into three distinct components - *Run, Report*, and *Refine* (Figure 1). We will discuss below the details of the *Run* pipeline, prioritized with accurate identification of gene loci and exon-intron boundaries. Both the *Report* and *Refine* pipelines accept the genes predicted in the *Run* pipeline in order to perform protein annotation and gene locus refinement, respectively, using the same overall parallelization techniques. The *Report* pipeline annotates proteins identified in the *Run* or *Refine* pipelines using established databases and tools such as MMseqs2 (Steinegger and Söding, 2017), KofamScan (Aramaki et al., 2019), and eggNOG (Huerta-Cepas et al.,2018), while the *Refine* pipeline accepts RNA-seq and/or transcriptome data that can be mapped to genomes using Hisat2 (Kim et al., 2019) and GMAP (Wu and Watanabe, 2005), respectively, and used in locus boundary refinement with BRAKER2 (Bruna et al., 2021). The implementation of the tools in the *Report* and *Refine* pipelines does not deviate from prescribed methodologies and accepts all user-defined parameters from the source programs.

**Figure 1:**
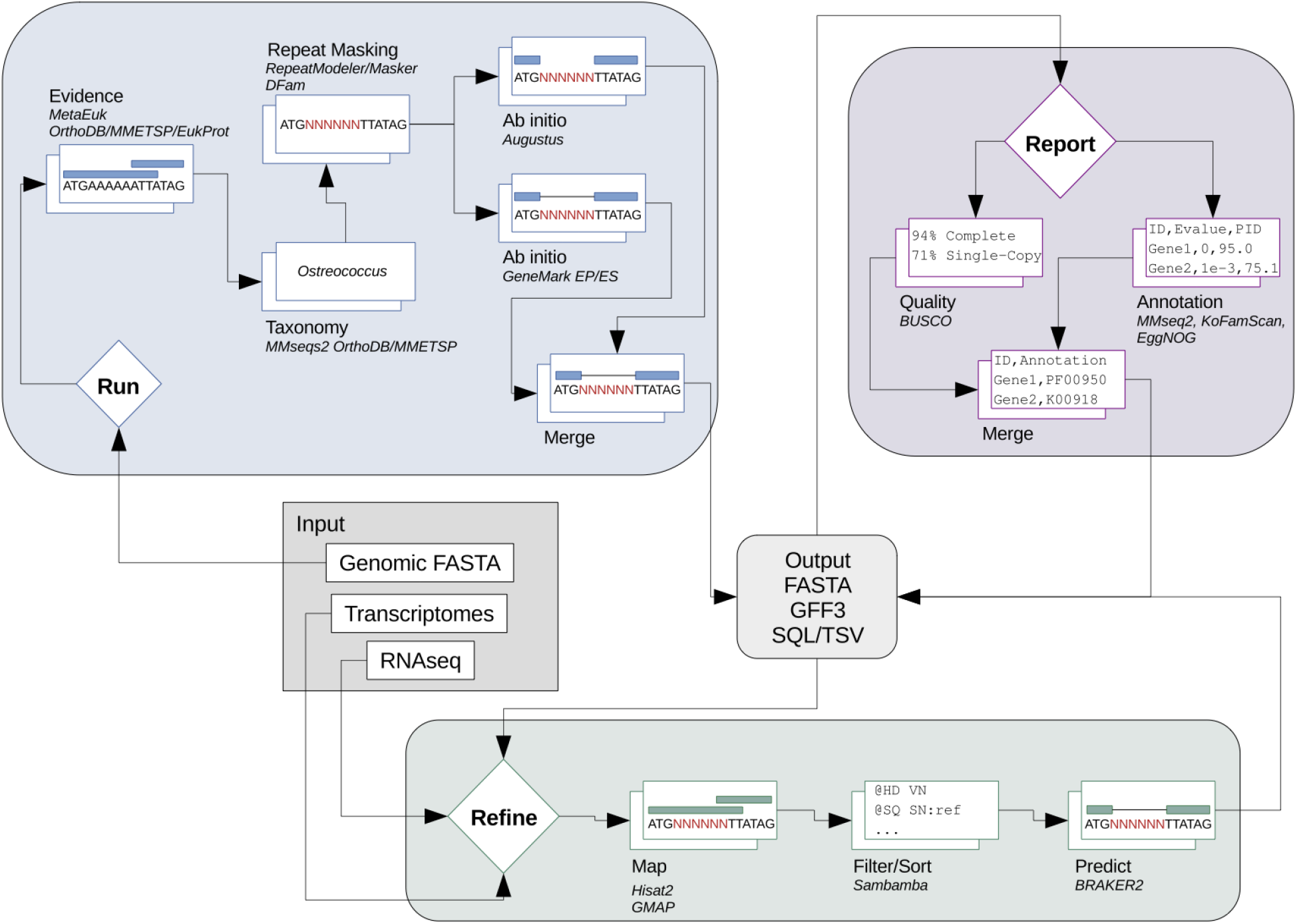
Schematic of the three EukMetaSanity pipelines: *Run, Report*, and *Refine*.

Herein, we will detail the performance of the *Run* pipeline, which automates gene locus prediction. The first priority of the *Run* pipeline is the determination of an approximate NCBI taxonomic assignment for the genome of interest, as this assignment will inform repeat masking and the proteins used as evidential support in gene prediction. This assignment is completed on a first-pass set of protein predictions that are generated by the program MetaEuk (Levy Karin et al., 2020). The MMseqs2 taxonomy subcommand compares the input genome against a modified database that contains the Orthologous Database of Proteins (OrthoDB; n = 1,271 eukaryotic genomes) (Kriventseva et al., 2019) and the Marine Microbial Eukaryotic Transcriptome Sequencing Project (MMETSP; n = 719 transcriptomes) (Keeling et al., 2014). The combined OrthoDB-MMETSP database provides extended coverage beyond laboratory cultivars and macrofauna to include environmental eukaryotes, specifically emphasizing marine protists. The combined dataset encompasses 1,482 species and provides representatives for 352 Orders in 127 Classes (overlap determined using https://github.com/frallain/NCBI_taxonomy_tree/pull/1). For reasons discussed below, these databases were selected due to their use of the NCBI taxonomy ID (taxid) ontology schema, which provides the ability to extract related organisms based on a shared identifier. Additional databases that use the NCBI taxid can be easily incorporated to expand the breadth of the databases packaged with EukMetaSanity, but databases that lack this shared ontology (Niang et al., 2020; Richter et al., 2020) would require modification in their corresponding taxonomy schema or integration into steps further downstream in the *Run* pipeline than currently implemented.

When an appropriate NCBI taxonomy can be identified, genomes undergo repeat identification and masking using RepeatMasker (Smit et al., 2013) which uses the Family- or Superfamily-level NCBI taxid to select repeat models from the DFam library (Hubley et al., 2016). Repeats are also masked in an *ab initio* fashion using RepeatModeler2 (Flynn et al., 2020), which runs multiple iterations of repeat identification and refinement to generate repeat families that inform the masking step in RepeatMasker. Taxonomic information is then used to select all relevant proteins from the OrthoDB-MMETSP database that match at least the Order-level predicted assignment. These proteins are used as inputs in gene prediction for GeneMark-EP (Bruna et al., 2020). Should a suitable Order-level assignment be unavailable or GeneMark ProtHint fail to predict intron boundaries, the annotation step defaults to GeneMark-ES (Lomsadze et al., 2005) to perform *ab initio* prediction. Additionally, we automate the first round of Augustus (Stanke et al., 2006) training by searching the input genome against the OrthoDB-MMETSP database with the MMseqs2 subprogram linsearch and, for the 60 models in the Augustus species database, select the model with the highest scoring linsearch match. Predicted gene loci from all three annotation tools (GeneMark-EP/ES, Augustus, MetaEuk) are directly available, but, additionally, EukMetaSanity can combine multiple annotation tracks to provide gene loci approximations that capture all non-overlapping recovered loci (Tier 1), gene loci supported by two tools (Tier 2), or only gene loci supported by all three tools (Tier 3; Figure S1).

To explore how reliable the EukMetaSanity *Run* pipeline was in returning accurate gene predictions, multiple experiments were conducted using high-quality genomes with accompanying gold standard annotations from the NCBI Reference Sequence (RefSeq) and GenBank databases, as well as performing comparisons between the methodologies used to predict genes in environmentally derived MAGs. To illustrate that the protein evidence methodology used by EukMetaSanity can recapitulate the gene content of gold standard eukaryote genomes, 102 genomes were selected from the NCBI RefSeq database plus an additional set of 10 genomes tested using BRAKER2 (Bruna et al., 2021) and the recently released platypus genome (Zhou et al., 2021). The 112 genomes were selected to provide representatives from a phylogenetically diverse set of organisms with a range of overall genomic complexity and size, ranging from unicellular algae (*Guillardia theta*, 55.1Mbp) (Curtis et al.,2012) to the platypus (*Ornithorhynchus anatinus*, 1.86Gbp; Supplemental Data 1).

To our knowledge, this is the first time that a comparison has been made for 100+ eukaryote genomes using these three annotation approaches, with previous assessments ranging from 7-12 genomes (Levy Karin et al., 2020; Bruna et al., 2021; Banerjee et al., 2021). This computationally intensive task (18,461 CPU hrs for 48 genomes with length 100-400 Mbp; 8,879 CPU hours for 14 genomes ≥400 Mbp) is achieved in relatively short time-scales through the aggressive use of parallelization and optimization by EukMetaSanity to manage the resources distributed to compute nodes on an HPC system (Figure S2; Supplemental Data 2).

For the three gene prediction software suites used to analyze the NCBI genome set, BUSCO (Seppey et al., 2019) completion scores were used to estimate the impact on genome annotation (Supplemental Data 1). Genomes annotated with the GeneMark-EP program (Bruna et al., 2020) produced a distribution of identified BUSCO completion estimates that were not significantly different from the reference set (*p*_BH-FDR_ = 0.3553; Wilcoxon rank-sum). When compared to the reference annotation, the GeneMark-EP annotation resulted in a median decrease of five identified BUSCO proteins, while the median decrease for identified BUSCO proteins for Augustus (Stanke et al., 2006) and MetaEuk Levy Karin et al. (2020) was 70 and 110, respectively (Figure 2A). GeneMark-EP and MetaEuk over-predicted the total number of proteins in the dataset by 16.4% and 72.2%, respectively, while Augustus under-predicted the total number of proteins by 36.2% (Figure 2A). Combining annotation tracks using the Tier 1 criteria resulted in a final gene track which benefited from the initial high-scoring GeneMark-EP annotation set and which incorporates additional gene loci identified either by MetaEuk or Augustus, resulting in a median decrease of four BUSCO proteins when compared to the reference and an over-prediction of 65.7% of the total number of proteins (Figure 2A). Tier 1 predictions resulted in the smallest number of genomes that had lower BUSCO completeness, and was able to return the same completeness score or higher for 50.0% of the genomes. Collectively, the GeneMark-EP and Tier 1 approach performed well in minimizing the number of BUSCO proteins lost during gene prediction; however, both methods over-predict the number of detected genes. The high degree of over prediction in the Tier 1 approach is to be expected as it combines results from all three tools, substantially increasing the number of false positives. As an alternative approach, the EukMetaSanity Tier 2 method outputs genes that are supported by at least two lines of evidence. With the Tier 2 criteria, the number of genomes that lost BUSCO proteins increased, but this loss of true positives is offset by a drastic decrease in potential false positives (Tier 2 under-predicts total proteins by 6.3%; Figure S3). The use of either GeneMark-EP, Tier 1, or Tier 2 outputs depends on the goal of the researcher to maximize sensitivity (Tier 1) or precision (Tier 2; Figure S3).

**Figure 2:**
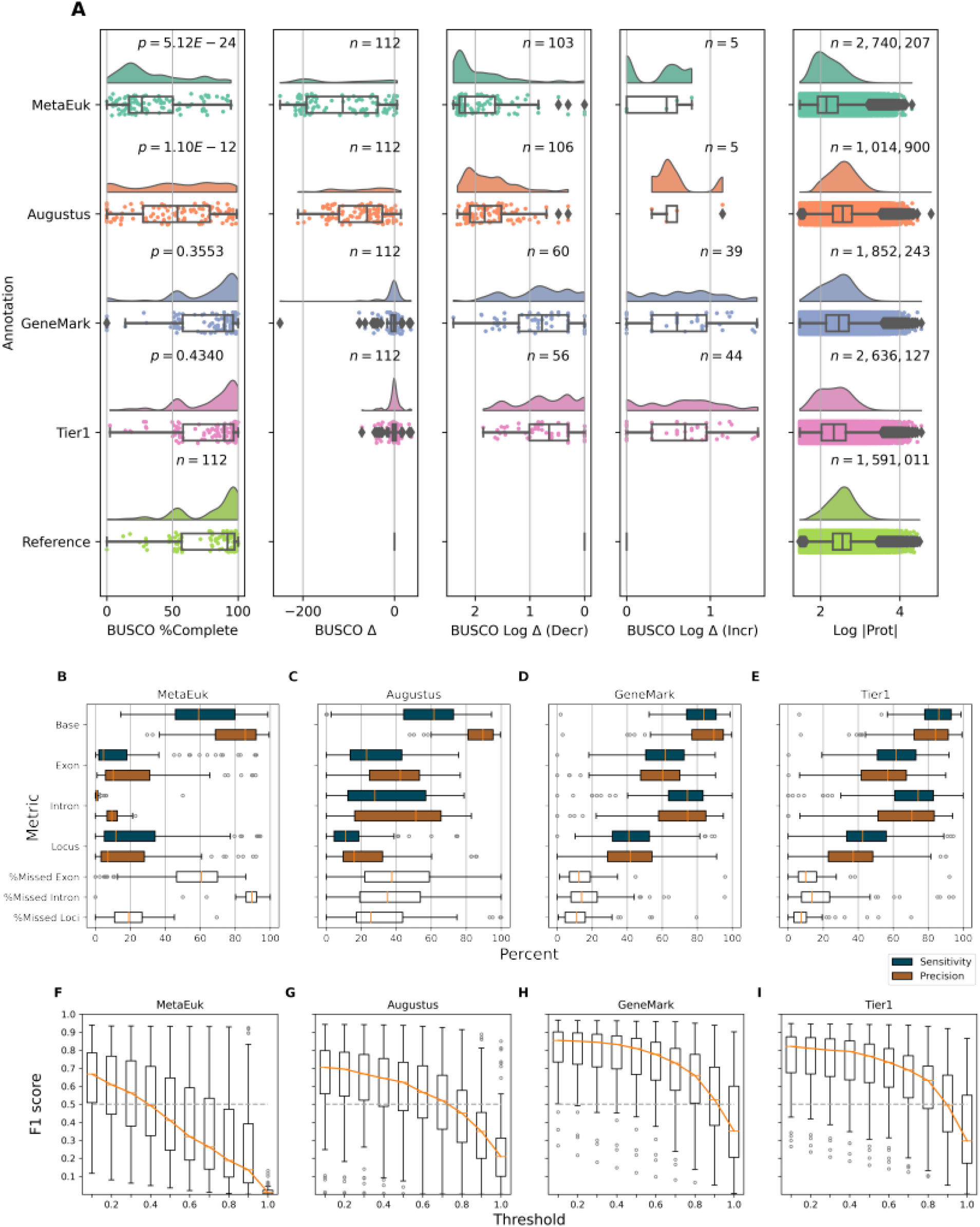
Comparison of EukMetaSanity results to NCBI provided annotations for 112 genomes. (A) Box plots comparing BUSCO completeness and protein prediction results for the three gene prediction tools and Tier 1 approach against the NCBI reference (*n* = 112). From left to right: Panel 1 - BUSCO completeness for each genome. *p*-value for Wilcoxon ranked sum with Benjamini-Hochberg false discovery correction. Panel 2 - The total number of BUSCO proteins lost/gained for each genome. Panel 3 - The log decrease in BUSCO proteins. Panel 4 - The log increase in BUSCO proteins. Panel 5 - Log size of proteins recovered. (B-E) Box plots comparing GffCompare results for the three gene prediction tools and Tier 1 approach. Sensitivity and precision calculated as *S* = *TP*/(*TP* + *FN*)) and *P* = *TP*/(*TP* + *FP*), respectively. %Missed Exon, Intron, and Locus indicates that value not recovered by the associated method. (F-I) Box plots comparing LocusCompare results for the three gene prediction tools and Tier 1 approach. *F*1 = 2 × (*S* × *P*)/(*S*+*P*)

To assess the recovery at the individual gene loci level, the program GffCompare (Pertea and Pertea,2020) was used to perform a stringent comparison between the EukMetaSanity output and the NCBI reference (Supplemental Data 1). GffCompare uses stringent cutoffs in its assignment of true positives (TP), false positives (FP), and false negatives (FN) that require exact coordinate matches for features (*i.e*., base-, exon-, intron-, and locus-level), and allows for no more than a 100-base difference at the ends of exons when assessing locus-level accuracy. We found that each of the programs used in EukMetaSanity achieved median sensitivity and precision values > 59.0% in the NCBI dataset when considering base-level matches (total number of bases assigned to an exon at the same coordinate), with the Tier 1 approach and GeneMark-EP scoring the highest base-level sensitivity (86.2% and 84.2%, respectively) and precision scores (83.8% and 89.5%, respectively; Figure 2B-E). Key differences in each program are apparent in the annotation quality with respect to accurate identification of exon and intron feature levels. While both features retained median sensitivity and precision > 59% for GeneMark-EP and Tier 1 predictions, MetaEuk saw median sensitivity and precision scores of 4.7% and 10.2%, respectively, for exon recovery, and 0.9% and 9.5%, respectively, for intron recovery (Figure 2B). For Augustus, the median sensitivity and precision were 23.1% and 42.3%, respectively, for exon recovery, and 27.8% and 51.2%, respectively, for intron recovery (Figure 2C). The median percentages for unannotated gene loci for MetaEuk, Augustus, GeneMark, and Tier 1 predictions were 19.2%, 25.5%, 11.1%, and 7.4%, respectively (Figure 2B-E). These results demonstrate that even when considering the strict cutoffs used by GffCompare, GeneMark-EP and the Tier 1 output are able to produce high accuracy annotation predictions that recover ~90% of the previously annotated loci.

At locus-level resolution, GffCompare results provide a perspective that relies on exact boundary accuracy of all introns. We performed a complimentary assessment to compare the degree to which gene content is captured, but for which intron structure may be incorrect. This method, which we call LocusCompare, identifies the degree to which a predicted gene overlaps a reference gene loci (*i.e*., length of prediction-reference overlap divided by the length of the reference; Figure S4). TP, FP, and FN are determined using a sliding scale of threshold cutoffs (0.1 - 1.0) that reflects the proportion of the reference locus that is recovered (*e.g*., a threshold value of 0.1 indicates a predicted gene overlaps at least 10% of the reference gene at a locus position and 1.0 represents an overlap of the exact length, or greater, of the reference gene; Figure S4; Supplemental Data 1). GeneMark-EP and Tier 1 predictions retained median F1 scores ≥ 0.5 for LocusCompare threshold values ≤ 0.9 (Figure 2F-G), while MetaEuk and Augustus F1 scores dropped < 0.5 at lower threshold values (≤ 0.4 and ≤ 0.7, respectively; Figure 2H-I; Supplemental Data 3). The GeneMark-EP and Tier 1 results indicate that, for many genes predicted by EukMetaSanity, the recovered genes are at least 90% of target gene length. And, while Tier 1 predicts a larger number of total proteins, this does not result in a large decrease in F1 scores when compared to GeneMark-EP. These results suggest that EukMetaSanity functions well when the status about the presence/absence of a protein is important, for example interpreting metabolic potential from a large MAG dataset (as in Alexander *et al*. 2021). A fraction of a loci, when translated to protein, may be suitable for detecting a gene despite inexact intron boundaries, and may yield results that can be explored further with the databases provided in the *Report* pipeline. We do not recommend the use of the EukMetaSanity *Run* pipeline alone if the goal is to determine exact intron-exon boundaries, which will still require additional transcript-level support.

From the larger overall analysis of 112 NCBI genomes, here we highlight 11 particularly relevant bellwether examples that underscore the advantages and limitations of EukMetaSanity for plant and animal taxa. For these taxa, EukMetaSanity was effectively able to recover a large percentage of the known gene loci (Figure S5). In this subset, the Tier 1 and Tier 2 approaches recovered BUSCO scores that differed by < 3.6% from each other, with the exception of the platypus genome. In these instances, the Tier 2 approach showed marked increase in F1 scores across all thresholds, as well as an increase in sensitivity and precision metrics at the base-, exon-, intron-, and locus-levels (Supplemental Figure 6F & 6K). The Tier 1 approach overestimated the number of recovered gene loci (760,818 predicted genes vs 384,839 reference genes), but the Tier 2 approach acted as a useful filter to remove false positives, pseudo-genes, and other spurious ORFs that are not supported by at least two annotation programs (349,810 predicted genes; Figure S5). The only exception to this was the *Ornithorhynchus anatinus* (platypus) genome, which exemplifies an instance when *ab initio* prediction is superior to protein evidence supported predictions. There are only 18,894 proteins in the OrthoDB-MMETSP from the Order *Monotremata*, which come from organisms other than *O. anatinus* (*i.e*., echidnas). In comparison, the platypus has 38,847 annotated genes. In this instance, *ab initio* prediction through Augustus was successful in recovering a large fraction of the expected genes (70.2% BUSCO completeness; Supplemental Data 1) compared to gene predictions that included protein evidence (7.5% and 0% BUSCO completeness for MetaEuk and GeneMark-EP, respectively). The lack of sufficient representation in the OrthoDB-MMETSP database drastically decreases the likelihood that protein evidence-based gene prediction software can accurately resolve gene loci in this newly sequenced mammal species.

To further explore the impact of a lack of closely related organisms in the taxonomic database on gene prediction results for novel environmental organisms, we artificially removed a selection of organisms from the OrthoDB-MMETSP database prior to gene prediction. From NCBI RefSeq, we identified 34 fungal genomes (assigned to the Kingdom Fungi) from 15 different Orders and created 15 modified OrthoDB-MMETSP databases, each depleted of proteins from the corresponding fungal Order (Supplemental Data 4). These results recapitulate those reported using the NCBI gold standard genomes: GeneMark-EP and the Tier 1 were superior in recovering predicted gene loci compared to Augustus and MetaEuk (Figure 3; Figures S6 and S7). Exploring the results from GeneMark-EP, there was no statistical difference between the BUSCO completeness for genomes when using the full database versus the database depleted of genomes sharing the same Order (*p*_BH-FDR_ = 0.849; Wilcoxon rank sum; Figure 3). For the GeneMark-EP output, we did see slight decreases in precision and sensitivity for most of the categories assessed using GffCompare and slight increases in the percent exons, introns, and loci missed in the annotation step. The total number of proteins recovered increased when using the depleted database (+42,850 proteins), while the average protein length decreased (−7 amino acids), suggesting that at least some of the gene loci were being unintentionally split when using a more distantly related protein set as evidence (Table S1). On an approach-by-approach basis, there was a trend of similar BUSCO completeness values and LocusCompare F1 scores when comparing the full versus depleted databases, though the values for MetaEuk and Augustus were substantially lower (Figure 3; Figure S7; Supplemental Data 5). The results from this assessment indicate that the method implemented by EukMetaSanity can be used to provide gene annotations for genomes even when near neighbors within the same Order are not available. In this instance, the presence of related Orders in the database acted as a bridge to provide sufficient information to recover quality gene annotations. For organisms that lack representation at the Class or Phylum level, the lack of reference proteins will undoubtedly have an impact on gene recovery. EukMetaSanity is designed to handle these cases by defaulting to the GeneMark-ES implementation and providing an output like the Tier 2 approach which can return putative gene loci supported by at least two of provided annotation tools, affording added confidence in the final annotation set, as well as decreasing the total false positive protein predictions that are included (Figure S3). As the data illustrates, this approach is also useful for more complex organisms.

**Figure 3:**
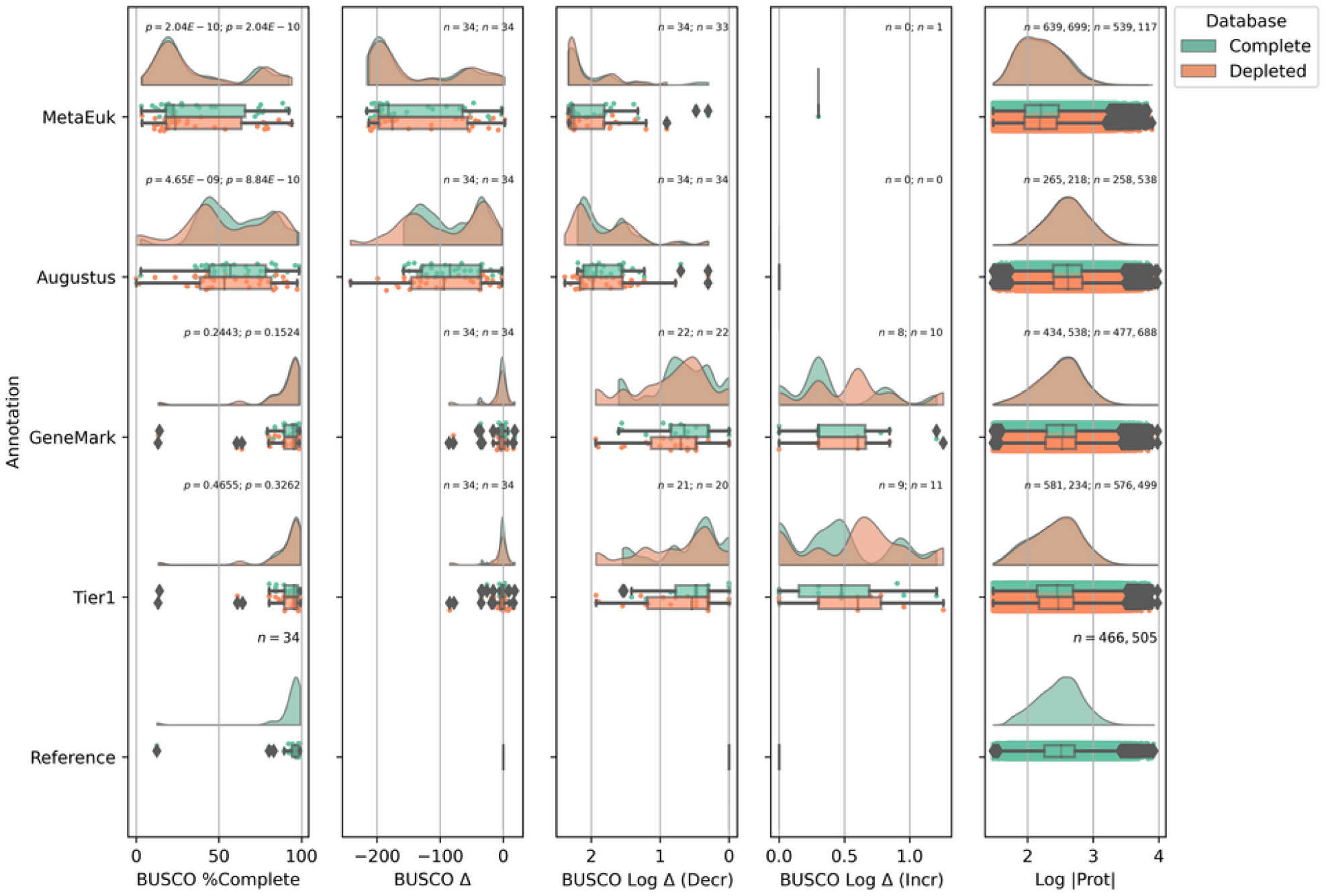
Reliability of EukMetaSanity using Order-level depleted databases for 34 fungal genomes in 15 Orders. Box plots comparing BUSCO completeness and protein prediction results for the three gene prediction tools and Tier 1 approach against the NCBI reference (*n* = 34) for the complete OrthoDB-MMETSP database (green) and depleted database lacking the associated Order-level set of proteins (orange). From left to right: Panel 1 - BUSCO completeness for each genome. *p*-value for Wilcoxon ranked sum with Benjamini-Hochberg false discovery correction. Panel 2 - The total number of BUSCO proteins lost/gained for each genome. Panel 3 - The log decrease in BUSCO proteins. Panel 4 - The log increase in BUSCO proteins. Panel 5 - Log size of proteins recovered.

After establishing the functionality of EukMetaSanity on the benchmark datasets above, we applied the *Run* pipeline to two sets of marine, eukaryotic MAGs reconstructed from the *Tara* Oceans large size fraction metagenomic datasets (Supplemental Data 6-7) (Carradec et al., 2018). Here we highlight the impact of applying EukMetaSanity gene prediction compared to two different approaches used by the authors. As no “gold standard” annotation exists for the recovered MAGs, our intentions were to compare the results provided by EukMetaSanity to the methodologies initially used in gene annotation.

Delmont *et al*. (2021) reconstructed 682 eukaryotic MAGs and performed annotation using a protocol that included protein mapping against the Uniref90 and METdb (Niang et al., 2020) databases with splice aware mapping, *ab initio* predictions using Augustus, and a mapping step that used 905 metatranscriptome assemblies (Pesant et al., 2015). While the specific approaches to protein mapping differ, the overall concept was conserved between the Delmont *et al*. (2021) methodology and the approach implemented by EukMetaSanity (*i.e*., providing useful protein evidence to downstream processes). However, the metatranscriptome mapping is a time and computationally intensive step (682 MAGs × 905 metatranscriptomes), where the total *Tara* Oceans metatranscriptome dataset totals ~ 11TB of data. Additionally, no repeat modeling or masking was conducted as part of the initial protocol. Within EukMetaSanity, GeneMark-EP and the Tier 1 results generated BUSCO completion estimates that were not significantly different from the completion estimates from the Delmont *et al*. (2021) protocol (*p*_BH-FDR_ = 0.2421; Wilcoxon rank-sum; Figure 4A). The Tier 1 approach increased the number of identified BUSCO proteins in over 62% of the processed MAGs by a median of five additional BUSCO proteins per MAG. For the less than 48% of MAGs that saw a decrease in BUSCO proteins, the median number of BUSCO proteins lost was three (Figure 4A). The Tier 1 approach increased the total number of identified proteins in the complete dataset by 19% (comparatively, GeneMark-EP identified 6% more proteins; Figure 4A). Both MetaEuk and Augustus under-predicted the number of genes in the MAG dataset and produced BUSCO completion estimates with values that were significantly lower than the Delmont *et al*. (2021) protocol (*p*_BH-FDR_ = 3.48 × 10^−10^, *p*_BH-FDR_ = 1.98 × 10^−5^, respectively; Wilcoxon rank-sum; Figure 4A). The fact that EukMetaSanity delivers similar MAG completeness scores without the computationally intensive transcriptome mapping, while increasing the total number of putative proteins, demonstrates that EukMetaSanity can be used to provide initial gene predictions for environmental genomes. Excitingly, when a dataset like the *Tara* Oceans metatranscriptome is available, the *Refine* pipeline is available to automate this step and provide refined gene predictions through BRAKER2.

**Figure 4:**
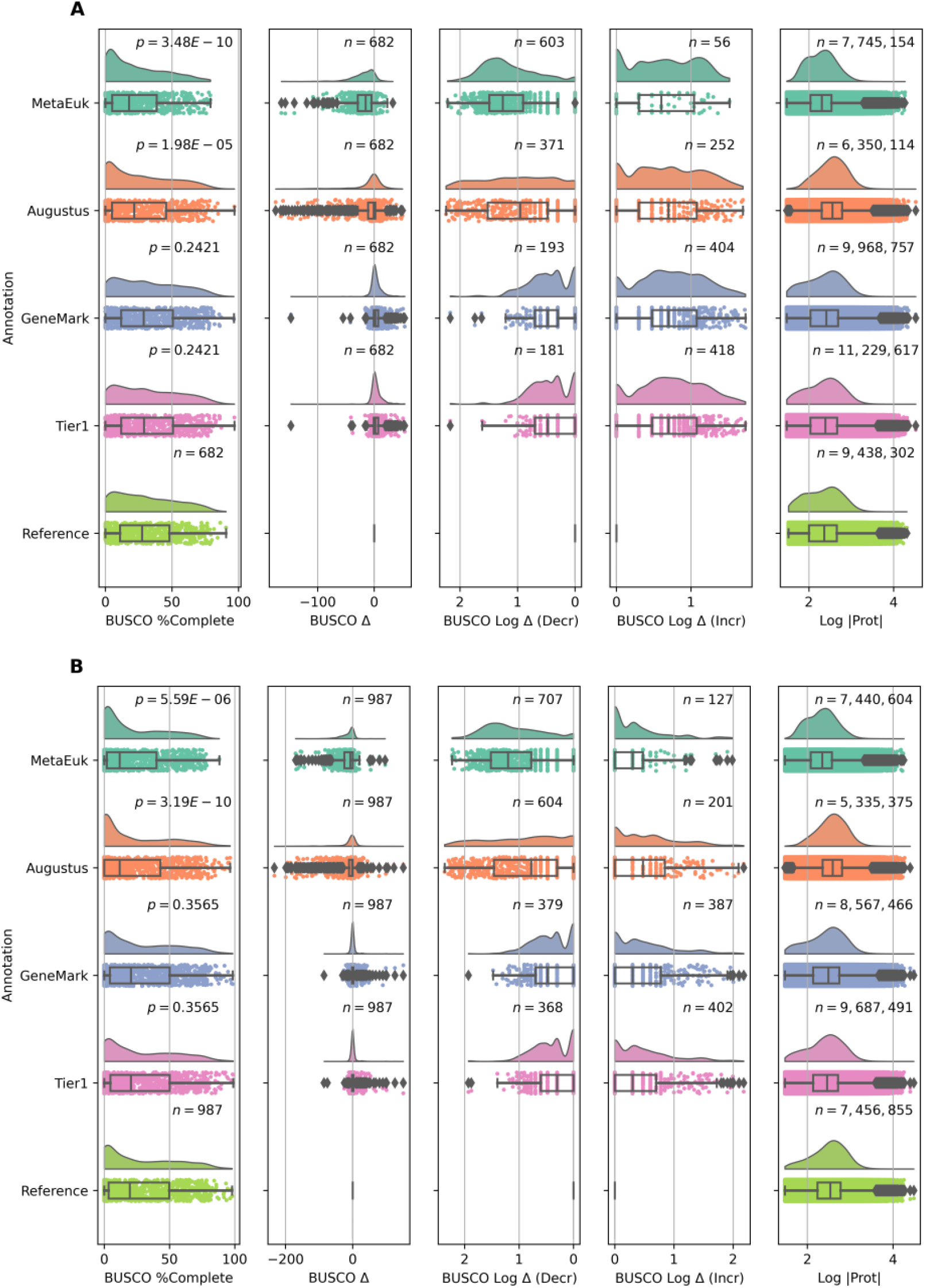
Comparison of EukMetaSanity results to alternative annotation pipelines used for *Tara* Oceans MAGs. (A) MAGs from Delmont *et al*. (2021) (n =682). (B) MAGs from Alexander *et al*. (2021) (n =987). From left to right: Panel 1 - BUSCO completeness for each genome. *p*-value for Wilcoxon ranked sum with Benjamini-Hochberg false discovery correction. Panel 2 - The total number of BUSCO proteins lost/gained for each genome. Panel 3 - The log decrease for genomes that lost BUSCO proteins. Panel 4 - The log increase for genomes that gained BUSCO proteins. Panel 5 - Log size of proteins recovered.

Interestingly, one putative Delmont *et al*. (2021) MAG (TARA_MED_95_MAG_00445) consistently failed to annotate genes with GeneMark-EP. Using genes identified with Augustus and MetaEuk (16 and 1,930 proteins, respectively), annotation with EukMetaSanity resulted in 1,943 non-overlapping protein predictions. Taxonomy prediction using the MMseqs2 taxonomy subprogram for the MetaEuk-derived proteins identified 44% as belonging to the Phylum *Ciliophora* and the MAG had a relatively small proportion (0.33%) of interspersed repeat elements (*n* = 322). A cursory analysis using Tiara (Karlicki et al., 2021) revealed that the 13.5 Mbp MAG in question consisted of DNA sequences whose origins were 28.8% eukaryotic, 28.3% bacterial, 10.2% archaeal, 7.3% prokaryotic, and 25.4% unknown. When protein prediction was performed using Prodigal v2.6.3 (-p meta) (Hyatt et al.,2012), a tool for prokaryotic gene prediction, the number of recovered putative coding sequences increased from 1,943 to 9,020, suggesting that this particular MAG represented a binning error that combined genomic content across Domains. This is an interesting test case to illustrate that the correctly implemented gene annotation pipeline can act as a quality control check on environmentally derived genomes going forward.

Alexander *et al*. (2021) generated 987 eukaryotic MAGs from the *Tara* Oceans large size fraction metagenomic dataset (Supplemental Data 7). The initial annotation protocol (not published) applied an *ab initio* pipeline that ran GeneMark-ES and MAKER (Holt and Yandell, 2011) on the input MAG with no masking of repetitive DNA. Using this annotation as a baseline, both GeneMark-EP and the Tier 1 approach were not significantly different from the estimated BUSCO completeness scores compared to the original Alexander *et al*. (2021) protocol (*p*_BH-FDR_ = 0.3565; Wilcoxon rank-sum; Figure 4B). Both MetaEuk and Augustus had significantly lower BUSCO completeness (*p*_BH-FDR_ = 5.59 × 10^−6^, *p*_BH-FDR_ = 3.19 × 10^−10^, respectively; Figure 4B). The Tier 1 approach increased the number of identified BUSCO proteins in 402 MAGs (40.7%) with 60 MAGs (6.1%) seeing an increase of ≥ 10 BUSCO proteins, including four MAGs (0.4%) seeing an increase of ≥ 100 BUSCO proteins. This approach maintained the number of identified BUSCO proteins in 217 MAGs (22.0%; Figure 4B). Conversely, 368 MAGs (37.3%) decreased in the number of identified BUSCO proteins, but only 18 MAGs (1.8%) lost ≥10 BUSCO proteins (Figure 4B). Using the Tier 1 approach, we increased the total number of proteins identified by 30%. Processing the Alexander *et al*. (2021) MAGs dataset illustrates how EukMetaSanity increases the quality of gene prediction and the total number of proteins from such a dataset, profoundly expanding the amount of data that can be passed to the functional annotation step and used for resolving metabolic reconstructions. The corresponding EukMetaSanity-derived gene predictions were used to assess functional potential for the Alexander *et al*. MAGs and provided insight into the ecological distribution of trophic strategies for the dataset (Alexander et al., 2021).

EukMetaSanity is an advanced workflow package for high-throughput gene prediction of eukaryotic genomes that streamlines the recovery of gene loci in environmental and cultivated genomes. Implementing the EukMetaSanity *Run* pipeline, we were able to annotate 1,785 genomes with repeats, introns, and exons using multiple top-of-the-line tools with automated selection for the necessary training data to recover high-quality gene loci information. The work presented here demonstrates that using the Tier approaches or GeneMark-EP as implemented within EukMetaSanity fits an important niche for the near future of eukaryotic genome annotation - both rapid preliminary annotation of genomes lacking transcript evidence and annotation of environmental genomes reconstructed from undersampled and uncultivated clades. When processed serially, many of the individual steps implemented in EukMetaSanity can be incredibly time intensive (e.g., >30 hours for repeat prediction and >24 hours for gene loci prediction for >1Gbp genomes) which can create bottlenecks during data processing. The automated parallelization provided by EukMetaSanity allows researchers to fully optimize their computational infrastructure, prevent wasted allocation allotments, and, because each step of EukMetaSanity is compartmentalized, provide an avenue of rapid parameterization of steps for variations in datasets. Rapid parameterization is especially important when new tools or databases are introduced into the workflow - which the API has been specifically designed to accommodate by making large-scale changes accessible with minimal understanding of the core code. EukMetaSanity provides a first step towards making the annotation of eukaryote genomes as accessible to the average researcher as current methodologies allow for their prokaryotic counterparts.

## Methods

EukMetaSanity was implemented using Python and the yapim API (https://github.com/cjneely10/YAPIM). Briefly, the yapim API distributes a data analysis pipeline across multiple inputs using user-defined memory and CPU resources. By leveraging simple concurrent computing operations such as locks and conditions, yapim reduces the total execution time for a resource-intensive analysis such as EukMetaSanity, allowing for and automating the parallelized large-scale analyses of hundreds or thousands of organisms.

### Databases

EukMetaSanity is packaged with three pre-computed MMseqs2 protein databases (Steinegger and Söding, 2017) which are used throughout each pipeline as a source of protein-level evidence to various programs. The databases are the Orthologous Database of Proteins (OrthoDB; n = 1,271 eukaryotic genomes) (Kriventseva et al., 2019), the Marine Microbial Eukaryotic Transcriptome Sequencing Project (MMETSP) (Keeling et al., 2014) protein taxonomy database (n =719 transcriptomes), and a combined version of the two (OrthoDB-MMETSP) generated by using the concatdbs subcommand of the MMseqs2 software suite. These databases contain embedded NCBI taxonomy information which affords sub-setting at various taxonomic levels. Several downstream tools are constrained by the use of NCBI taxonomy and require synchronization between databases at multiple phases, providing one of the main limitations between mixing tools and databases from multiple sources. By default, EukMetaSanity incorporates the OrthoDB-MMETSP database at various steps in its pipelines to provide adequate coverage of varying biologically-relevant datasets and environments. Users have the option of selecting to use any of the three provided databases, or to use their own MMseqs2 seqTaxDB database type.

Tools within EukMetaSanity rely on protein evidence to generate accurate predictions. The OrthoDB-MMETSP database is used to identify putative taxonomy and to generate a subset of proteins that are mapped to the input genome for intron boundary prediction. The choice of protein database will ultimately affect the kinds of evidence that are available in the pipeline. For example, the MMETSP database underrepresents animals, so using it to annotate a Metazoan genome without including OrthoDB can drastically affect the quality of an annotation.

### Run

The *Run* pipeline predicts gene loci and exon-intron boundaries for submitted genomes by accumulating *ab initio* and protein evidence. For each input genome, the *Run* pipeline will generate a set of one or more gene-finding format files (*e.g*., GFF, repeats), as well as FASTA files containing the gene coding regions of the genomic DNA and the derived protein sequences. The results from each program are optionally merged to a single set of non-overlapping gene locus predictions.

#### Protein evidence gene prediction

First pass protein prediction is completed using MetaEuk (Levy Karin et al., 2020) and the OrthoDB-MMETSP database. This step is completed on the input genome prior to masking, and identified proteins may include pseudo-genes that include larger repetitive DNA elements. MetaEuk predicts proteins by first performing a six-frame translation on each contig, identifying putative protein-coding segments. These segments are then searched through the OrthoDB-MMETSP database, and protein fragments that match to the same reference protein are collected. Fragments corresponding to putative exons are ordered and scored, and the highest scoring set of exons is returned as the putative protein. Here, users may provide e-value cutoffs and minimum sequence identities per the MetaEuk documentation.

#### Taxonomy

Subsequent steps in the *Run* pipeline rely on protein-based evidence at varying taxonomic levels. The MMseqs2 subprogram taxonomy provides an assignment of the MetaEuk-predicted protein sequences generated using OrthoDB-MMETSP. Users may provide e-value, coverage, and sequence identity cutoffs per the MMseqs2 documentation. In reported taxonomic assignments, levels that do not meet user-provided cutoffs are labelled with their parent labels (*i.e*., if the Order-level is not identified, pipeline steps will operate on the Class-level, etc.).

Next, sequences with identified homologs in the database are parsed into a taxon tree, and the NCBI taxid of the lowest common ancestor is identified between the query and matching protein sequence sets. The MMseqs2 taxonomyreport submodule converts the results of the taxonomy search to a taxon tree that displays the organism’s likely assignment at various taxonomic levels (Kingdom, Phylum, *etc*.). Users may provide a lower-bound on the acceptable percentage of proteins that need to be identified for an assignment.

#### Repeats identification

The input genome is processed in order to hard-mask short and long interspersed nuclear repeats, as well as other DNA transposons, small RNA, and satellite repeats. Two options for repeat identification have been included as part of the *Run* pipeline. RepeatMasker (Smit et al., 2013) identifies repetitive DNA content from the DFam library (Hubley et al., 2016) of repeats using either the Family- or Superfamily-level NCBI taxid identified in the taxonomy step. RepeatModeler2 (Flynn et al., 2020) can be used as the only source of repeat prediction or in conjunction with a RepeatMasker species to generate *ab initio* predictions of repetitive DNA regions. Including both levels of repeat annotation maximizes the chances to identify and mask repeat content in the genome. RepeatModeler has a long runtime and operates best on genomes with high assembly N50, and RepeatMasker requires that NCBI repeat models exist for a given genome. Users who do not perform both steps may miss repeat content. Because repetitive DNA regions may include pseudo-genes whose intron boundaries do not reflect intron models of true genes, these regions must be masked prior to subsequent *ab initio* prediction to generate the most accurate models of intron splice sites in a given organism. Both of these DNA repeat models are then used to hard-mask any repeat regions identified, excluding low-complexity repeats. Users may also provide external repeat libraries to use in masking at this step, if they are available or for more complex organisms.

#### *Ab initio* gene prediction

EukMetaSanity contains implementations for two *ab initio* gene prediction programs. In both cases, repeat-masked genome sequences are used as input. Users may use one or both *ab initio* protocols. Using both protocols affords the additional opportunity to capture gene content that may be missed by the other predicting software.

> Augustus (Stanke et al., 2006): Augustus is packaged with gene identification models for various animals, alveolates, plants/algae, fungi, bacteria, and archaea. EukMetaSanity automates the selection of an Augustus species model for each input genome by running the MMseqs2 linsearch module to identify proteins in the masked genome sequence that are found in the OrthoDB-MMETSP database. The Augustus species that bears the highest number of assignments to proteins identified by linsearch is selected as the model for the first round of *ab initio* training. Users may provide search criteria to linsearch that specify minimum sequence identity, coverage, *etc*.
>
> Subsequent rounds of training are conducted iteratively by generating training parameters based on the results of the prior round. This is accomplished using the Augustus subprograms gff2gbSmallDNA.pl, new_species.pl, and etraining. Typically, one additional round of Augustus training is sufficient, but users may elect to perform more training rounds as desired. High numbers of training rounds may result in a gradual reduction in captured gene content, so users are advised against over-training.
>
> GeneMark-EP/ES (Bruna et al., 2020; Lomsadze et al., 2005): Using the MMseqs2 module filtertaxseqdb, proteins that share the same Order-level assignment as the input genome are subset from the OrthoDB-MMETSP database. These proteins are provided as input to the Gene-Mark subprogram ProtHint, which generates intron predictions by performing spliced protein alignments of the sequences selected from OrthoDB-MMETSP to the masked genome using the program Spaln (Gotoh, 2008). If ProtHint fails, or < 100 introns are predicted, then proteins are predicted using GeneMark-ES. Otherwise, output from ProtHint is provided with the masked genome as input to GeneMark-EP. Users may provide parameters to either ProtHint or GeneMark-EP to filter the allowable contig and gene sizes in the prediction protocol. GeneMark-EP or -ES is automatically run in fungal mode for relevant genomes.

#### Merging final results

While the output of each preceding program can be used directly, we provide a final merging protocol to reduce the gene annotations to a single set of gene predictions per gene locus in poly-logarithmic time (https://github.com/cjneely10/LocusSolver). Each annotation file is input into the merging script in GFF3 format. Annotation locations (loci) are merged into larger “superloci” which can consist of one or more gene tracks at that locus. For each strand within each superlocus, a predicted gene is selected, with priority selection assigned to GeneMark-EP/ES, then Augustus, and finally to MetaEuk (Tier 1; Figure S1). Users may add an additional filter to retain only genes whose locus is supported by more than one line of evidence (Tier 2 or Tier 3; Figure S1).

### Refine

The *Refine* pipeline is a transcriptome-based gene prediction workflow that uses BRAKER2 (Bruna et al., 2021) for predictions. The repeat-masked genome and Order-level proteins subset from the OrthoDB-MMETSP database generated in the *Run* pipeline are used as inputs. Users may also choose to incorporate the gene prediction results from a previously completed GeneMark-EP/ES implementation in the *Run* pipeline, or may start a new *ab initio* gene prediction process. Users will provide either trimmed RNA-seq reads or assembled transcriptomes as input to the pipeline. Trimmed RNA-seq data are mapped to the masked genome using Hisat2 (Kim et al., 2019), the SAM output of which is subsequently converted to BAM format and sorted using Sambamba (Tarasov et al., 2015). Assembled transcriptomes are mapped to the input genome using GMAP (Wu and Watanabe, 2005), the results of which are then similarly converted and sorted to BAM format. The combined set of output BAM files are provided as input to BRAKER2, which outputs multiple result formats including Augustus- and GeneMark-based predictions. The additional level of evidence provided by gene expression data captures feature content such as alternative transcription start sites and the locations of 5’ and 3’ untranslated regions of genes. Additionally, gene expression data can modify and improve results from *Run*-derived predictions.

### Report

The *Report* pipeline provides functional annotation of the predicted proteins. This pipeline annotates gene loci predicted in either the *Run* and/or *Refine* stages using MMseqs2 (Steinegger and Söding, 2017), eggNOG-emapper (Huerta-Cepas et al., 2019, 2018), and/or KofamScan (Aramaki et al.,2019). MMseqs2 runs the subprogram search or linsearch against a number of pre-computed databases provided from the MMseq2 authors (mmseqs databases command), including Pfam (El-Gebali et al., 2018), UniRef (Suzek et al., 2007), the NCBI non-redundant protein database, db-CAN2 (Zhang et al., 2018), and others, all of which can be used to assign functional annotation to genes. The download of these databases occurs outside of EukMetaSanity, but any number of pre-computed or user generated databases can be designated for search with this step within the configuration file. eggNOG-emapper and KofamScan are implemented within this pipeline to provide eggNOG and KEGG functional annotations to proteins, respectively. These programs are distributed through their specific organizations and must be installed externally with instructions provided on the EukMetaSanity installation page (https://github.com/cjneely10/EukMetaSanity/blob/main/INSTALLATION.md). Tab-delimited summary files for each gene call are generated.

### Benchmark Datasets

We tested how well EukMetaSanity accurately predicted proteins using three different datasets. Parameters used with EukMetaSanity can be found in Table S2. We collected a set of 112 gold standard genomes from NCBI (National Library of Medicine, US) from various taxonomic levels across the tree of life (accessed 7 January 2021; Supplemental Data 1). We selected 34 fungal genomes that belonged to 15 fungal Orders (Supplemental Data 4). For each Order, we generated an MMseqs2 database from the combined OrthoDB-MMETSP database that removed all proteins from the target Order using the MMseqs subprogram filtertaxseqdb. Each fungal genome was then annotated using the EukMetaSanity *Run* pipeline with a respective database depleted in proteins from their identified Order. EukMetaSanity was tested against a large eukaryotic MAG dataset of 1,669 MAGs, generated from the *Tara* Oceans metagenomic datasets by Delmont *et al*. (2021) (n = 682 MAGs; Supplemental Data 6) and Alexander *et al*. (2021) (n = 987 MAGs; Supplemental Data 7). In Delmont *et al*. (2021), the MAGs were annotated using a tripartite approach that integrated protein alignments, metatranscriptomic mapping, and *ab initio* gene predictions. In Alexander *et al*. (2021), the MAGs were annotated using GeneMark-ES and the MAKER pipeline with no repeat masking. Both sets of MAGs were annotated using the EukMetaSanity *Run* pipeline with the complete OrthoDB-MMETSP databases.

### Benchmarking

We assessed gene prediction performance at the whole genome level by comparing BUSCO (Seppey et al., 2019) completeness results with the eukaryota_odb10 dataset, and the sensitivity and precision of individual gene loci for EukMetaSanity results compared to the annotations provided by NCBI. Each of the four gene loci prediction protocols (MetaEuk, GeneMark-EP, Augustus, and the Tier 1 approach) were compared against the NCBI reference annotations.

For all gene predictions from NCBI references, MAGs, and EukMetaSanity, protein-level annotations in GFF/GFF3 format were converted to protein sequences using GFFread v0.12.1 and subsequently processed by BUSCO v4.1.2 (parameters: -m prot -l eukaryota) to provide a completeness estimate for each genome. We compared the distribution of BUSCO completeness scores identified in the reference annotations to the EukMetaSanity annotations from each gene prediction protocol with significance determined using the Wilcoxon rank-sum statistic (Mann and Whitney, 1947) with Benjamini-Hochberg multiple test correction (Benjamini and Hochberg, 1995).

Using the program GffCompare v0.12.5 (Pertea and Pertea, 2020), we generated metrics that compared the reference annotations to annotations derived from the programs contained within EukMetaSanity. These metrics summarized the sensitivity (*S* =*TP*/(*TP* + *FN*)) and precision (*P* = *TP*/(*TP* + *FP*)) between the reference and query annotations at the base, exon, intron, and intron-chain (locus) levels. At the base-, exon-, and intron-levels, a query feature is only marked as a true positive if the start and end coordinates exactly match the reference. For intron chains, all introns in the query transcript must exactly match the reference transcript. At the locus level, a putative intron chain in the query locus must exactly match an intron chain in the set of intron chains present at the reference locus. Intron chain-level accuracy allows a maximum distance of 100bp between the ends of the terminal exons in the query locus and the ends of the reference locus.

We developed LocusCompare (https://github.com/cjneely10/EukMetaSanity/blob/main/tests/LocusCompare.py) as an additional metric that determines gene-specific sensitivity and precision based on an overlapping locus in the genome and compares the predicted gene to the reference gene (Figure S4). We determined the fraction of the reference gene covered by the prediction by dividing the length of the overlapping region by the total length of the reference gene. We then set threshold cutoffs values between 0.1 and 1.0 and considered features that meet the threshold as TPs, and those that do not meet the threshold (or that bear redundancies) as FPs. Any missed reference gene locus is marked as a FN. F1 scores were calculated for each annotation set for each threshold (*F*1 = 2 × (*S* × *P*)/(*S* +*P*)).

## Supporting information

Supplmental Informtaion

## Data and Code Availability

Supplemental Data 1-7 and the EukMetaSanity-generated gene, protein, and repeat predictions for all NCBI reference genomes and MAGs are hosted through figshare. Supplemental Data 1-7: https://doi.org/10.6084/m9.figshare.15044334; NCBI genomes: https://doi.org/10.6084/m9.figshare.15040554; fungal genomes with depleted databases: https://doi.org/10.6084/m9.figshare.15042633; Delmont *et al*. (2021): https://doi.org/10.6084/m9.figshare.15042645; Alexander *et al*. (2021): https://doi.org/10.6084/m9.figshare.15042636. EukMetaSanity and accompanying code is available at https://github.com/cjneely10/EukMetaSanity. Euk-MetaSanity is licensed through the GNU General Public License v3.0. And yapim, the API used to construct the steps in EukMetaSanity, can be found at https://github.com/cjneely10/YAPIM.yapim is licensed through the Creative Commons Attribution-Non Commercial 4.0 International License.

## Acknowledgements

The authors would like to thank the staff that support their respective HPC systems: Poseidon at the Woods Hole Oceanographic Institution, Discovery at the University of Southern California Center for Advanced Research Computing, and Endeavor in the USC Quantitative and Computational Biology Department. The authors would also like to thank the *Tara* Oceans consortium for providing open access to their expansive “omics” datasets and other researchers who embrace rapid and open access to their data. SKH was supported by a Postdoctoral Fellowship from the Center for Dark Energy Biosphere Investigations (C-DEBI) through NSF-OCE-0939564 and by NSF-OCE-1947776. HA was supported by NSF-OCE-1948025 and a WHOI Independent Research and Development award. BJT was supported by C-DEBI through NSF-OCE-0939654. This is C-DEBI Contribution XXX.

## Author contributions statement

B.J.T., H.A., and C.J.N. conceived of and designed the study; C.J.N. wrote the code, performed the analyses, and analyzed the data; H.A. performed additional tests; B.J.T. and C.J.N. wrote the manuscript; S.K.H. provided expertise on database construction and usage; S.K.H. and H.A. provided edits to the manuscript. All authors reviewed the manuscript.

## Ethics Declaration

The authors declare no conflicts of interest.

