## Supplementary material for "The high-throughput gene prediction of more than 1,700 eukaryote genomes using the software package EukMetaSanity": Supplmental Informtaion

### Supplementary Figures and Tables for "The high-throughput gene prediction of more than 1,700 eukaryote genomes using the software package EukMetaSanity"

Christopher J. Neely[1,*], Sarah K. Hu[2], Alexander Harriet[3], and Benjamin J. Tully[4,5,*]

[1]University of Southern California, Department of Quantitative and Computation Biology, Los Angeles, 90089, USA
[2]Marine Chemistry and Geochemistry, Woods Hole Oceanographic Institution, Woods Hole, MA, USA, 02543
[3]Biology Department, Woods Hole Oceanographic Institution, Woods Hole, MA, USA, 02543
[4]University of Southern California, Wrigley Institute for Environmental Studies, Los Angeles, 90089, USA
[5]University of Southern California, Center for Dark Energy Biosphere Investigations, Los Angeles, 90089, USA
[*]

July 25, 2021

#### Supplementary Figures

Includes Figures S1-S7.

#### Supplementary Tables

Includes Tables S1-S2.

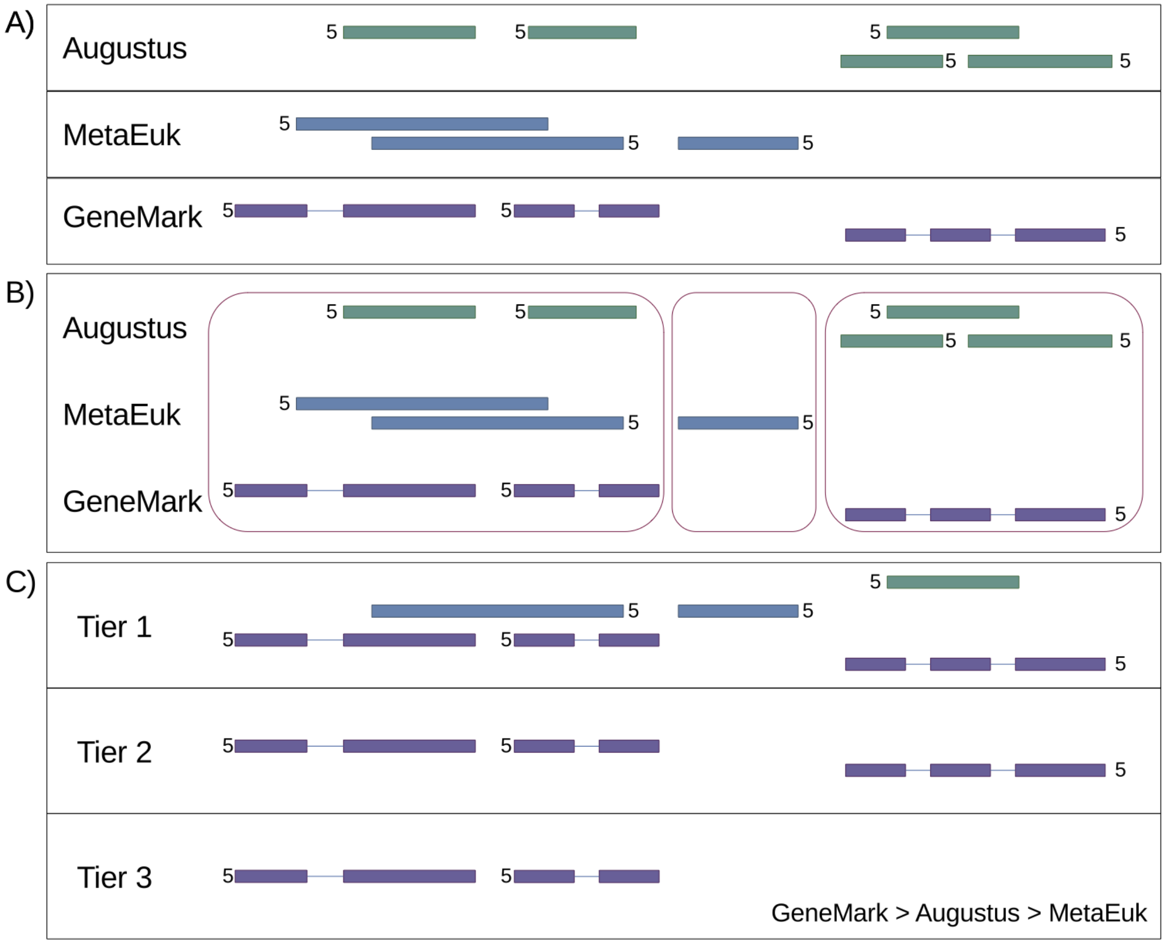


Figure S1: **Expectations for the Tier approach implemented in EukMetaSanity.** (A) Illustrates three output gene tracks from Augustus, MetaEuk, and GeneMark-EP/ES. (B) All predicted exon and intron segments that cover the same locus coordinates are considered together. (C) Expected predicted genes based on the Tier approach. When conflicting exon/intron boundaries occur in the same direction, outputs from specific tools are given priority: GeneMark-EP/ES > Augustus > MetaEuk. Tier 1 outputs all non-conflicting, detected gene loci. Tier 2 requires support from at least two gene prediction tools. Tier 3 requires support from all three tools.

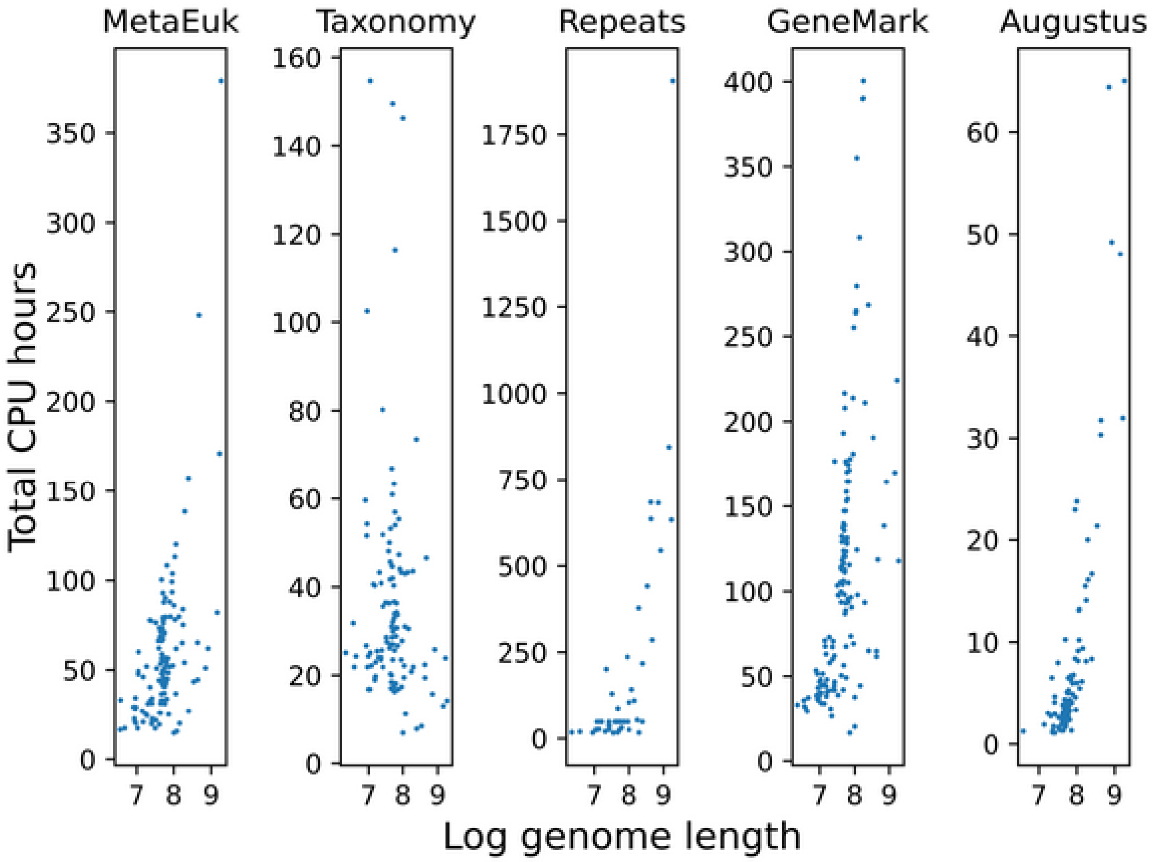


Figure S2: **Elapsed CPU time for other major steps in EukMetaSanity workflow.** Time in Repeats reflects complete time to mask. GeneMark represents genomes that may have used GeneMark-EP or defaulted to GeneMark-ES.

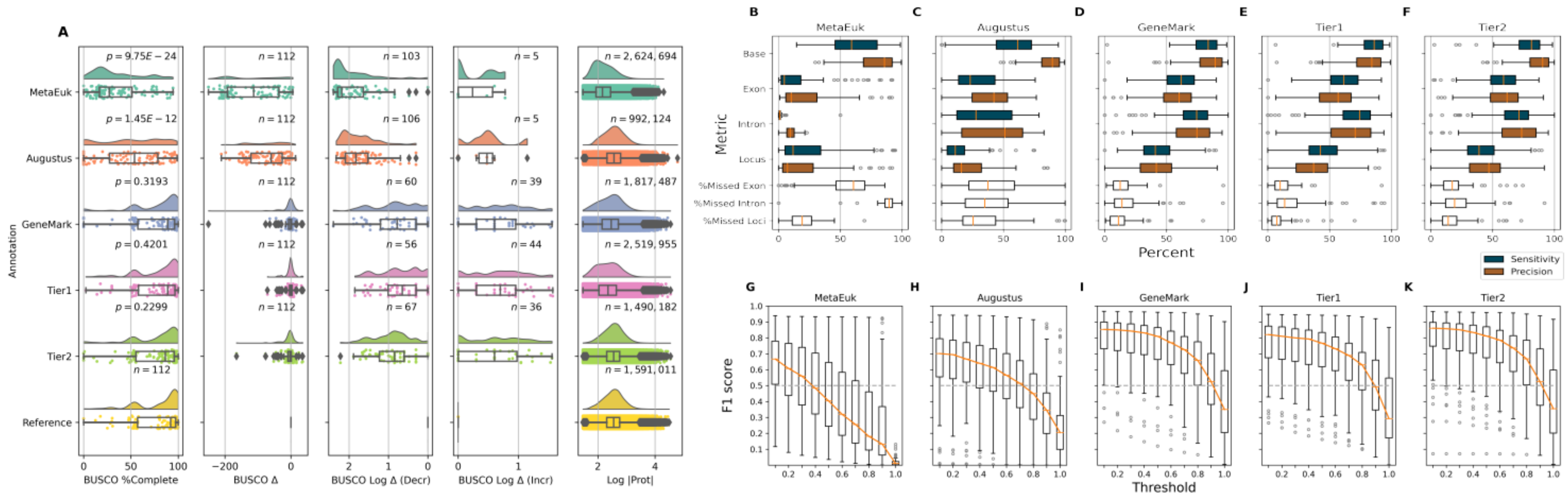


Figure S3: **Comparison of EukMetaSanity results to NCBI provided annotations for 112 genomes.** (A) Box plots comparing BUSCO completeness and protein prediction results for the three gene prediction tools, Tier 1, and Tier 2 approaches against the NCBI reference ($n = 112$). From left to right: Panel 1 - BUSCO completeness for each genome. $p$-value for Wilcoxon ranked sum with Benjamini-Hochberg false discovery correction. Panel 2 - The total number of BUSCO proteins lost/gained for each genome. Panel 3 - The log decrease for genomes that lost BUSCO proteins. Panel 4 - The log increase for genomes that gained BUSCO proteins. Panel 5 - Log size of proteins recovered. (B-F) Box plots comparing GffCompare results for the three gene prediction tools, Tier 1, and Tier 2 approaches. Sensitivity and precision calculated as $S = TP/(TP+FN)$ and $P = TP/(TP+FP)$, respectively. %Missed Exon, Intron, and Locus indicates that value not recovered by the indicated method. (G-K) Box plots comparing LocusCompare results for the three gene prediction tools, Tier 1, and Tier 2 approaches. $F1 = 2 \times (S \times P)/(S+P)$

.

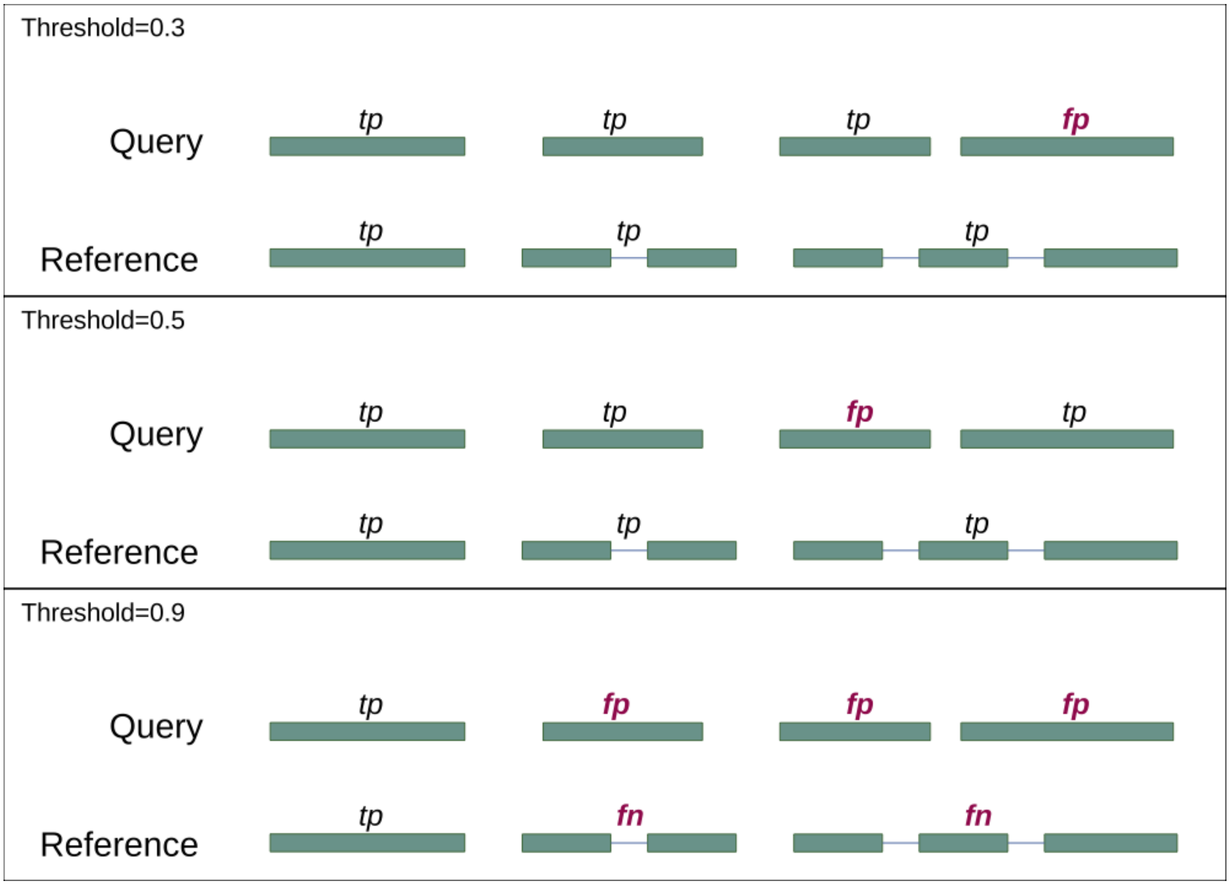


Figure S4: **Visualization as to how changes in the threshold score impact LocusCompare.** $Threshold = 0.3$, a query splits exons of one gene into multiple genes. Only one is considered if above the threshold length and all others are considered FP. For $threshold = 0.5$, the smaller predicted gene is not 50% of the length of the reference gene, but the previously identified FP meets this length threshold. Abbrev. tp, true positive; fp, false positive; fn, false negative.

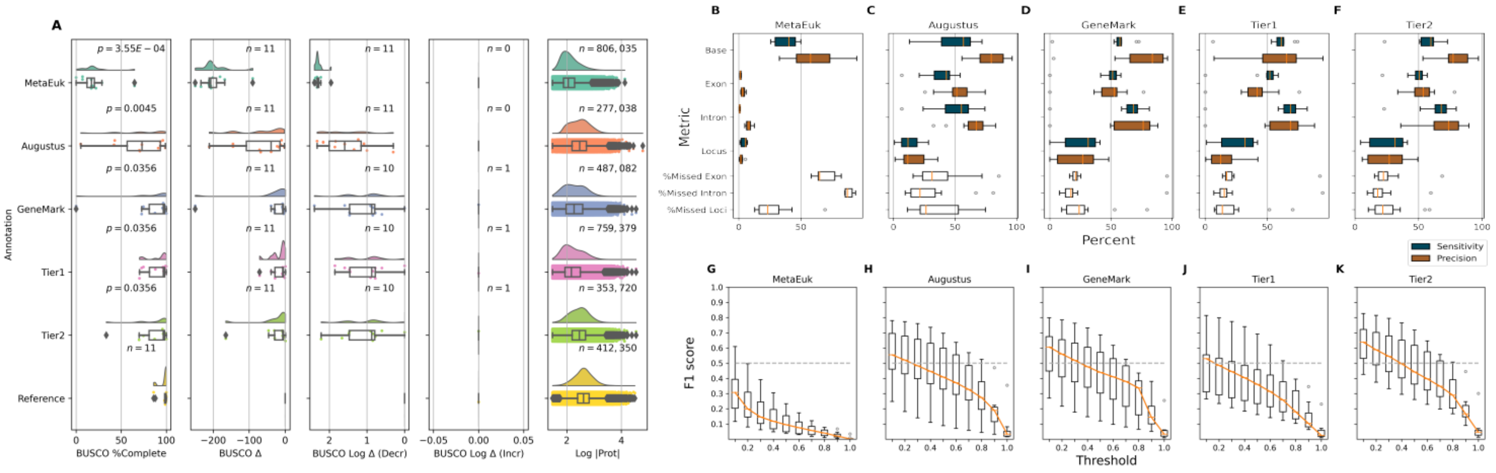


Figure S5: **Comparison of EukMetaSanity results to NCBI provided annotations for 11 plant and animal genomes.** Genomes in this subgroup include: *Arabidopsis thaliana, Caenorhabditis elegans, Drosophila melanogaster, Populus trichocarpa, Medicago truncatula, Solanum lycopersicum, Bombus terrestris, Parasteatoda tepidariorum, Tetraodon nigroviridis, Danio rerio,* and *Ornithorhynchus anatinus*. (A) Box plots comparing BUSCO completeness and protein prediction results for the three gene prediction tools, Tier 1, and Tier 2 approaches against the NCBI reference ($n = 11$). From left to right: Panel 1 - BUSCO completeness for each genome. $p$-value for Wilcoxon ranked sum with Benjamini-Hochberg false discovery correction. Panel 2 - The total number of BUSCO proteins lost/gained for each genome. Panel 3 - The log decrease in BUSCO proteins. Panel 4 - The log increase in BUSCO proteins. Panel 5 - Log size of proteins recovered. (B-F) Box plots comparing GffCompare results for the three gene prediction tools, Tier 1, and Tier 2 approaches. Sensitivity and precision calculated as $S = TP/(TP+FN)$ and $P = TP/(TP+FP)$, respectively. %Missed Exon, Intron, and Locus indicates that value not recovered by the indicated method. (G-K) Box plots comparing LocusCompare results for the three gene prediction tools, Tier 1, and Tier 2 approaches. $F1 = 2 \times (S \times P)/(S+P)$

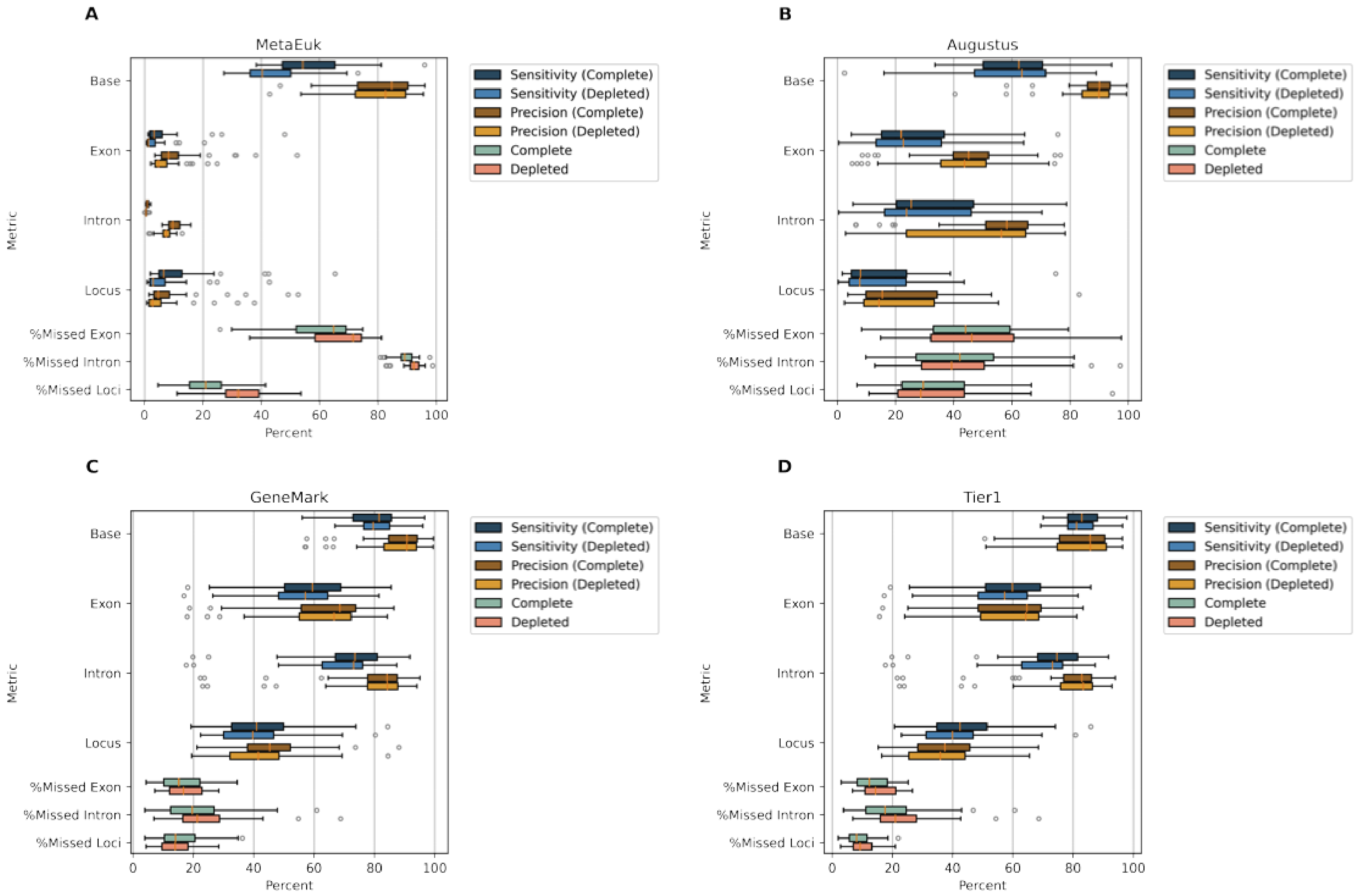


Figure S6: **Box plots comparing GffCompare results for the three gene prediction tools and Tier 1 approach.** Values are presented for both the complete ODB-MMETSP database (blue and aqua shades) and the depleted databases lacking the associated Order-level set of proteins (brown and coral shades). Sensitivity and precision calculated as $S = TP/(TP+FN)$ and $P = TP/(TP+FP)$, respectively. %Missed Exon, Intron, and Locus indicates that value not recovered by the associated method.

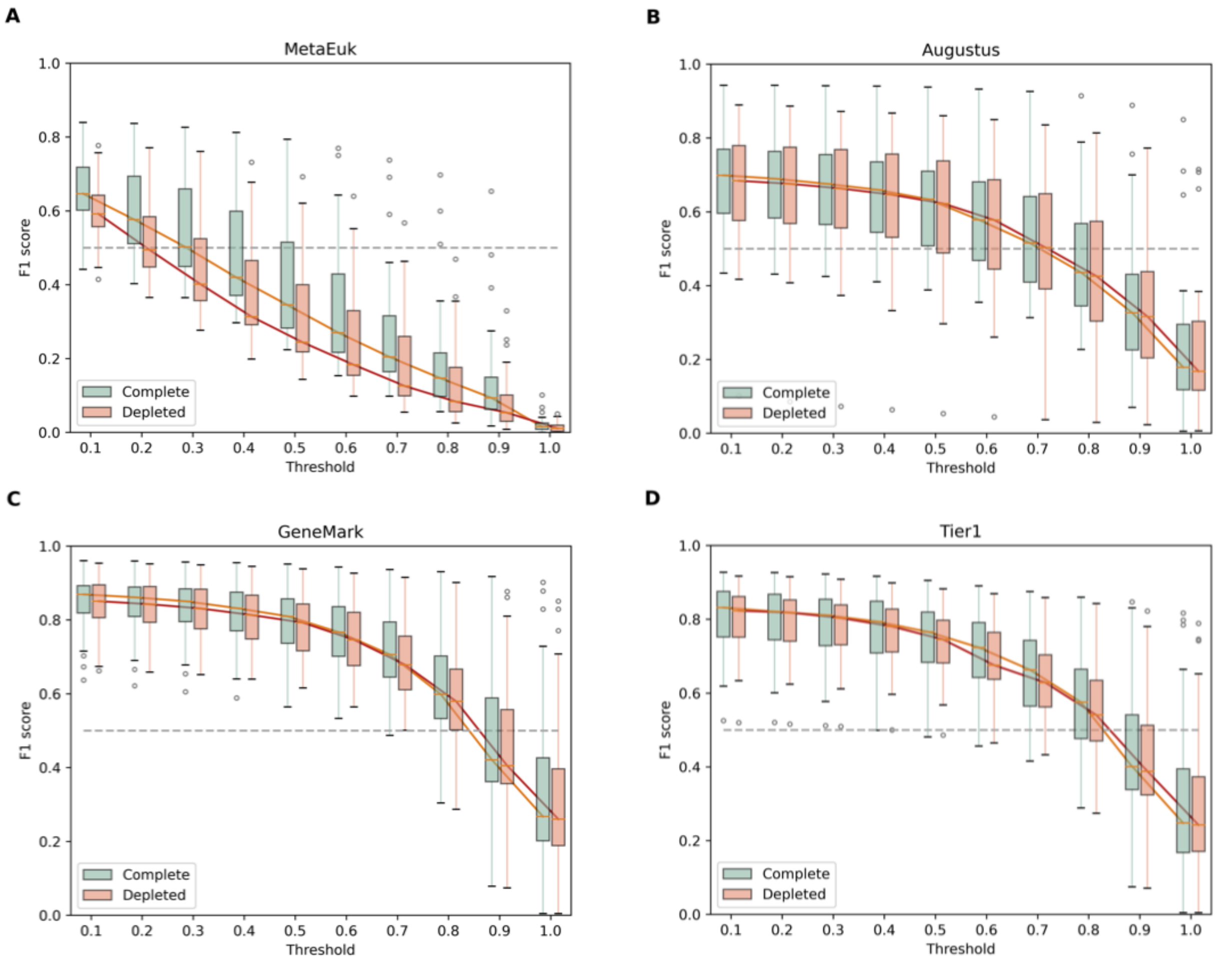


Figure S7: **Box plots comparing LocusCompare results for the three gene prediction tools and Tier 1 approach.** Complete ODB-MMETSP database (aqua) and the depleted databases lacking the associated Order-level set of proteins (coral). $F1 = 2 \times (S \times P)/(S + P)$.

Table S1: **Protein recovery values for the complete ODB-MMETSP database vs the Order-level depleted databases.**

| Protein values | MetaEuk complete | MetaEuk deplete | Augustus complete | Augustus deplete | GeneMark complete | GeneMark deplete | Tier 1 complete | Tier 1 deplete |
|---|---|---|---|---|---|---|---|---|
| **Count** | 639,699 | 539,117 | 265,218 | 258,538 | 434,538 | 477,388 | 581,234 | 576,499 |
| **Mean Length** | 169 | 162 | 404 | 413 | 321 | 314 | 260 | 272 |
| **Max length** | 6,831 | 7,875 | 9,541 | 9,541 | 9,525 | 9,529 | 9,525 | 9,529 |
| **Min length** | 30 | 30 | 30 | 30 | 30 | 30 | 30 | 30 |
| **1Q** | 91 | 89 | 245 | 249 | 194 | 186 | 139 | 149 |
| **Median** | 159 | 154 | 404 | 413 | 344 | 336 | 282 | 293 |
| **3Q** | 296 | 279 | 662 | 680 | 547 | 541 | 490 | 502 |

Table S2: **EukMetaSanity tasks, associated programs, and parameters.** Only parameters that deviate from those used for the NCBI reference genomes are noted.

| EukMetaSanity Task Name | Program Name | Citation | Version | NCBI parameters | Depleted DB parameters | Delmont parameters | Alexander parameters |
|---|---|---|---|---|---|---|---|
| **MetaEukEV** | MetaEuk | Karin<br>*et al.* 2020 | 4-a0f584d | --min-length 30<br>--metaeuk-eval 0.0001<br>-s 6 --cov-mode 0<br>-c 0.3 -e 100<br>--max-overlap 0 | -- | -- | -- |
| **Taxonomy** | MMSeqs<br>`taxonomy` | Steinegger<br>& Soding 2017 | 12.113e3 | -s 6.5 -c 0.3<br>--cov-mode 0<br>--min-seq-id 0.40 | -- | -- | -- |
| **Repeats** | RepeatMasker<br>ProcessRepeats | Smit<br>2005 | open-4.0.9 | -nolow | -- | -- | -- |
| **AbinitioGeneMark** | ProtHint | Bruna<br>*et al.* 2020 | 2.6.0 | --min_contig 500<br>--max_contig<br>5000000<br>--min_contig_in_predict<br>500<br>--min_gene_in_predict<br>30 | -- | -- | - |
| **AbinitioGeneMark** | PETAP | Lomsadze<br>*et al.* 2005 | 4.65_lic | --min_contig 500<br>--max_contig<br>1000000000000<br>--min_contig_in_predict<br>500<br>--min_gene_in_predict<br>30 | -- | --max_contig<br>100000000 | --max_contig<br>100000000 |
| **AbinitioAugustus** | MMSeqs<br>`linsearch` | Keller<br>*et al.* 2011 | 12.113e3 | --cov-mode 0<br>-c 0.6 -e 0.01 | -- | -- | -- |
